# Evaluation of the *Escherichia coli* population in the intestinal microbiota of rattus wistar submitted to secondhand smoke and supplemented with prebiotics

**DOI:** 10.1101/2020.03.25.008755

**Authors:** Bruna Rafaela dos Santos Silva, Paula Marioto Perez, Hermann Bremer Neto, Rogeria Keller

**Affiliations:** Universidade Estadual de Campinas, Instituto de Biologia, Departamento de Genética, Evolução e Bioagentes, Campinas, SP, Brazil; Universidade do Oeste Paulista, Departamento de Microbiologia, Presidente Prudente, SP, Brazil

**Keywords:** Microbiota, prebiotics, smoke, *E. coli*

## Abstract

Many chronic conditions, including smoking, have been shown to be associated with modifications in the gut microbiota and to remedy the effects of these changes, functional foods such as prebiotics have shown beneficial effects. The aim of this work was to evaluate changes in the bacterial population of *Escherichia coli* in the intestinal microbiota of rats submitted to passive smoke and to the supplementation of prebiotics incorporated into the basal diet. The animals were divided into: Control Group (CG) = basal diet; Smoking Control Group (GCT) = basal diet with exposure to smoking; Prebiotic Group (GP) = basal diet incorporated with 1g of Immunowall® per kg of diet; Prebiotic Tobacco Group (GPT) = basal diet incorporated with 1g of ImmunowallTM per kg of diet with exposure to smoking. The animals were exposed to the smoke twice a day for 180 days. The obtained data were analyzed through the nonparametric Kruklla-Wallis test. Bacterial populations were amplified by real-time PCR. The results of this study revealed a significant decrease (p = 0.04 and p = 0.01) in *Escherichia coli* population in the group of animals supplemented with prebiotics in the intestinal microbiota of rats exposed and not exposed to cigarette smoke reinforcing the modulating effect of potential pathogens on the intestinal microbiota by functional foods.

## 1. Introduction

The gut microbiota is composed of 10^13^ to 10^14^ microorganisms that are responsible for physiological function such as digestion and metabolism, of which the majority are bacteria colonized from birth (Gill et al., 2006). Many chronic conditions are associated with modifications of gut microbiota such as inflammatory bowel disease and obesity (Clavel et al., 2014). Previously studies in humans reported the relation between smoking and microbiota changes (Nolan-Kenney et al., 2019), showing differences in the microbiome measured in saliva (Wu et al., 2016), upper gastrointestinal tract (Vogtmann et al., 2015) and bronchial wash (Einarsson et al., 2016) samples. In additional, previously results suggested that smoking should be included in the growing list of known factors that influence the composition of the intestinal microbiota (Biedermann et al., 2013) increasing the risk of infections by pathogens and/or opportunistic bacteria (Bagaitkar, Demuth, & Scott, 2008).

To remedy the effects of changing the microbiota functional foods such as prebiotics, have shown beneficial effects. Thus a prebiotic can be defined as a selectively fermented ingredient that results in specific changes in the composition and / or activity of the gastrointestinal microbiota, conferring benefit (s) on the health of the host (Guimarães et al., 2019).

The bacteria *Escherichia coli* is a gram negative bacillus lactose fermenter, becomes part of the intestinal mucosa when ingested in food. This genus inhabits the large intestine in symbiosis, establishing a mutual relationship with the host, where it produces vitamins K and the B complex that are absorbed by the intestinal epithelium, which in turn provides nutrients for bacterial metabolism. However, when *E. coli* leaves the intestinal tract it can cause serious infections. Being assigned the term opportunistic pathogen (Delmas, Dalmasso, & Bonnet, 2015; Sharma et al., 2014). Thus, *E. coli* can be an extremely aggressive pathogen when it comes to pathogenic pathotypes such as: *E. coli* enteroinvasora, *E. coli* enteropathogenic, *E. coli* enterohemorragica, *E. coli* enterotoxigenic, where they adhere and can invade the intestinal mucosa, causing serious consequences to the host, such as the destruction of intestinal villi (Shoaf, Mulvey, Armstrong, & Hutkins, 2006).

In this way, it is in the interest of the scientific community to seek viable alternatives that can reduce the impacts caused by passive smoking on the human body. Therefore, the present study quantified and checked changes in the population mass of bacteria *Escherichia coli* that inhabit the intestinal tract of *Rattus novergicus* of the *wistar* strain, submitted to secondhand smoke for 180 days after intake of prebiotics.

## 2. Material and Methods

### 2.1 Animal ethics

The study was conducted according to the ethical principles in animal research adopted by the Brazilian College of Animal Experimentation and was approved by the (UNOESTE) Ethics Committee for Animal Research (No. 2656).

### 2.2 Experimental design

Thirty male albino rats of the *wistar* strain were analyzed between 45-50 g of body weight kept in individual cages, under the same lighting conditions (12/12-hour light / dark cycle), with controlled temperature around 22°C (Merusse & Lapichik, 1996) and the solid and water diets were provided *ad libitum* during the experimental period that lasted 180 days, with five days of adaptation to the management, food and basal diet and 175 days to treatments. The animals were randomly divided into four groups: Control Group (CG) = basal diet; Smoking Control Group (GCT) = basal diet with exposure to smoking for one hour daily, divided into two half-hour periods; Prebiotic Group (GP) = basal diet incorporated with 1 g of Immunowall® Mananoligosaccharide-MOS prebiotic (ICC USA, Inc., Louisville, Kentucky - USA) per kg of the diet; Prebiotic Smoking Group (GPT) = basal diet incorporated with 1g of MOS per kg of the diet with exposure to smoking for one hour daily, divided into two half-hour periods.

The animals were challenged to passive smoke in an average concentration during the exposure periods of 350 ppm CO2 measured through a specific gas detector, model TxiPro®, from Biosystems. To carry out this protocol, two hermetically sealed chambers were used, both being 100 cm long, 40 cm wide and 40 cm high. Commercially purchased cigarettes were used.

### 2.3 DNA Isolation and Real-time PCR

Fecal DNA was extracted with the aid of the DNA Mini-Stool Kit (QIAGEN) according to the manufacturer’s instructions. The quantification of DNA was performed on a NanoDrop ND-1000 spectrophotometer, NanoDrop Technologies. The evaluation of the intestinal microbiota was performed through real-time PCR of the 16S RNA region corresponding to the *Escherichia coli* gene. The Real-time PCR was performed using the Microbial DNA qPCR Assays Kit (QIAGEN) according to the manufacturer’s recommendations. The reaction was used to measure the number of single copies of the 16S rRNA gene.

### 2.3 Data analysis

To compare the results of smokers and non-smokers within the treatment, the Mann-Whitney non-parametric test was used. To compare between treatments within each group, the Kruskal-Wallis non-parametric test was used. The analyzes were conducted using the Biostat 5.3 software, considering 5% as the level of significance (Ayres et al., 2007).

## 3. Results and Discussion

Many chronic conditions, including smoking, have been shown to be associated with modifications in the gut microbiota and to remedy the effects of these changes, functional foods such as prebiotics, have shown beneficial effects. In this context, we conducted this work to compare the gut bacterial population of *E. coli* between groups submitted to passive smoking and supplementation with prebiotics and groups non-smoking and non-supplemented.

The results of the statistical analyzes are summarized in **Table 1** with the values of Cycle threshold (CT). The CT value corresponds to the point at which the threshold crosses the amplification line, allowing to identify the number of minimum cycles necessary for the amplification of the sequence of interest, being inversely proportional to the amount of DNA present in the sample (Nascimento, Suarez, & Pinhal, 2010).

**Table 1.** Cycle Threshold value for smoking and non-smoking animals (values expressed as mean ± standard deviation)

| Groups | *E.coli* population |  | P value |
| --- | --- | --- | --- |
|  | Control | Prebiotic |  |
| Smoking | 32.10 ± 1.13 | 33.81 ± 1.20 | 0.04* |
| Non-smoking | 23.09 ± 15.48 | 34.74 ± 0.71 | 0.01* |

p = value of statistical significance in the Kruskal-Wallis test; \* value of statistical significance in the Mann-Whitney test.

Our results revealed a significant decrease (p <0.05) with the administration of prebiotics of the *Escherichia coli* population in the intestinal microbiota of exposed and unexposed rats compared to the control group. The behavior of *E. coli* in the presence of prebiotics has been previously described (González-Ortiz et al., 2014; Molist et al., 2014; Tran, Everaert, & Bindelle, 2018), through receptors similar to those expressed by the host cells, where the microorganism binds to the surface of the prebiotic fiber by saturating its receptors, this way, it does not adhere to the intestinal mucosa, thus being excreted. Furthermore, the combined action of prebiotics and the probiotic *Lactobacillus rhamnosus* demonstrated to cause greater inhibition of a pathogenic *E. coli* striper in co-cultures analyzed (Anand, Mandal, & Tomar, 2019).

The existence of molecular mimicry between some sites of the prebiotic fiber and the receptors of the host’s intestinal epithelial cells was previously suggested, to which pathogenic microorganisms recognize and adhere, thus, these oligosaccharides act by adsorbing the pathogenic microorganisms to the intestinal epithelium. This interaction is explained by the presence of numerous receptors in pathogenic *E. coli* pathotypes that recognize in addition to the host cells, also the oligosaccharides (Moon, Whipp, Argenzio, Levine, & Giannella, 1983).

The fact that the reduction in the population of *Escherichia coli* is beneficial to the host, is based on the existence of pathogenic *E. coli* pathotypes, such as enterotoxigenic *E. coli*, enteropathogenic *E. coli*, enteroinvasive *E. coli*, enterohemorrhagic *E. coli*, being responsible for very serious intestinal infections, which express adhesins related to the pathogenesis of these bacteria and which are not expressed on the surface of commensals *E. coli* (Tran et al., 2018).

Thus, the statistically significant decrease in *E. coli* population in the intestinal microbiota of rats that received prebiotics and especially in rats exposed to cigarette smoke reflects the beneficial effect of prebiotics including in the microbiota with cigarette interference, where through prebiotic food damages caused by cigarettes can be modulated.

## 4. Conclusion

With the present study, we concluded that the administration of prebiotics caused a significant decrease in the population of *Escherichia coli* in the microbiota of rats exposed and not exposed to cigarette smoke, suggesting its modulating effect of the intestinal microbiota, even in a microbiota modified by cigarette smoke.

## Acknowledgements

The authors are grateful for the financial support provided by UNOESTE (proc. n°2656) and Fundação de Amparo à Pesquisa do Estado de São Paulo (FAPESP 2015/24261-3).

